# A Framework for Better Sensor-Based Beehive Health Monitoring

**DOI:** 10.1101/2022.11.15.516676

**Authors:** Asaduz Zaman, Alan Dorin

## Abstract

Hive bees provide essential pollination services to human agriculture. Managed honey bees in particular pollinate many crops, but also create honey and other bee products that are now of global economic importance. Key aspects of honey bee behaviour can be understood by observing hives. Hence, the limitations of manual observation are increasingly being addressed by new technologies that automate and extend the reach of hive monitoring.

Here we propose a framework to classify and clarify the potential for sensor-assisted hive monitoring to inform apiculture and, ultimately, improve hive bee management. This framework considers hive monitoring approaches across three newly proposed categories: Operational monitoring, Investigative monitoring, and Predictive monitoring. These categories constitute a new “OIP Framework” of hive monitoring. Each category has its own requirements for underlying technology that includes sensors and ICT resources we outline. Each category is associated with particular outcomes and benefits for apiculture and hive health monitoring detailed here. Application of these three classes of sensor-assisted hive monitoring can simplify understanding and improve best-practice management of hive bees.

Our survey and classification of hive monitoring to date show that it is seldom practiced beyond honey bees, despite the need to understand bumble bees and stingless bees also. Perhaps unsurprisingly, sensor-based hive monitoring is shown to remain primarily a practice of developed nations. Yet we show how all countries, especially developing nations, stand to gain substantially from the benefits improved sensor-based hive monitoring offers. These include a better understanding of environmental change, an increased ability to manage pollination, an ability to respond rapidly to hive health issues such as pests and pathogens, and even an ability to react quickly to the danger posed to insects and humans alike by extreme events such as floods and fires. Finally, we anticipate that the future of hive monitoring lies in the application of Predictive monitoring, such that a hive’s anticipated future state can be preemptively managed by beekeepers working iteratively with novel hive monitoring technologies.

## 1 Introduction

Approximately 87.5% of flowering plants [Ollerton et al., 2011] and, consequently 35% of global food products for human consumption, depend on effective pollination. Bees are key pollinators and provide essential ecosystem services to our natural and agricultural ecosystems [Hung et al., 2018]. Bee pollination can also improve the quality, shelf-life, and economic value of food [Klatt et al., 2014]. Honey, an abundant byproduct of some wild and domesticated social bees, has also been prized by humankind for thousands of years for its nutritional and medicinal value [Bogdanov et al., 2008]. Recently it has become an important global economic commodity [FAO, 2021]. Hive bees also have social and cultural significance worldwide [Prendergast et al., 2021, Perichon et al., 2020, Zamudio and Hilgert, 2018]. As a result of their value to humankind, understanding and supporting bees has long been, and remains today, of great importance. A fundamental way to improve our knowledge of bees is to actively monitor their behaviour and health, both in the wild and within the hives of managed colonies. This has, arguably, never been more important than today when native and wild pollinator insects face serious anthropogenic obstacles, including pesticide use [Gill et al., 2012, Kluser et al., 2010], agricultural intensification [Gill et al., 2012, Klein et al., 2007], habitat loss [Goulson et al., 2015, Potts et al., 2010, Kluser et al., 2010], climate change [Goulson et al., 2015, Potts et al., 2010] and extreme weather events [Goulson et al., 2015].

Monitoring social bees inside and outside their hives beyond research laboratories is valuable to understand their behaviour [Marchal et al., 2019]. However, close monitoring of beehives is not always straightforward. Difficulties can be introduced by the small size and sophisticated internal architecture of hives, as well as by the complex social behaviour of the inhabitants. These issues mean that studies dependent only on human observations have intrinsic limitations. For instance, it is quite infeasible for a human observer to track the individual behaviour of one, let alone many, of tens of thousands of tiny, nearly identical unmarked, fast-moving, free-flying insects that go about their daily activities both within the confines of a complex hive [Wario et al., 2017], and across landscapes many kilometres square [Ratnayake et al., 2021b]. Humans have only so much time, patience and observational capability. Monitoring selected properties of a hive in its entirety is certainly more manageable. This has been attempted many times previously [Meikle and Holst, 2015], and is the approach that underlies the bee hive monitoring framework we describe in this article.

The recent introduction of electronic and digital sensor technologies to bee monitoring has improved our understanding of the behaviour of social bees. For instance, research using temperature sensors has determined that changes in honey bee colony brood temperature can indicate a colony’s readiness to swarm [Zacepins et al., 2016]. Another study pinpointed the exact location inside the hive where this temperature change is most prominent [Zhu et al., 2019]. A third found evidence that vibrations within a colony are related to its brood cycle [Bencsik et al., 2015]. The use of sensor technology within and around hives, can be preferable to having human researchers poking, prodding, opening and closing hives, as it reduces colony disturbance while monitoring. It also simplifies the conduct of continuous, detailed, long-term studies.

Sensor technology’s falling costs, increasing sophistication, and ease of use, have lowered the barriers to its application beyond research. This enables beekeepers, even hobbyists, to adopt sensor-assisted hive monitoring. The commercialisation of hive sensors and monitoring systems has subsequently been undertaken. In light of this increasingly broad adoption, as with any system engaging diverse stakeholders, a structured framework will be beneficial to understand the needs of users, and the benefits they might expect from engagement. We did not presently find any such structure described in the literature. Therefore, this study proposes a new framework for understanding sensor-assisted beehive monitoring systems. Our intention is to facilitate technological adoption and adaptation, and direct innovation towards improvements in hive health monitoring. This is, as we show below, a key concern of current monitoring efforts.

It is not our intention to provide a systematic literature review of all hive monitoring technologies. Meikle and Holst (2015) [Meikle and Holst, 2015] have thoroughly reviewed different applications for continuous beehive monitoring and discussed sensor technologies. Marchal et al. (2019) [Marchal et al., 2019] have also briefly discussed sensors in the context of computational apidology. Our scope in this study is different. We have focused on monitoring approaches applied to bee species used as managed pollinators or for honey production (i.e. honey bees (Apidae: Apini), bumble bees (Apidae: Bombini), and stingless bees (Apidae: Meliponini)) to provide a detailed analysis and classification framework for existing beehive sensor technologies to guide future innovation. Our framework enables hive monitor users and developers to clearly articulate the kinds of monitoring they wish to conduct or support. This simplifies users’ and designers’ decision-making when selecting the technology required to support their goals. It can even inform decisions about whether or not technology is appropriate in particular cases. Sometimes, we are at pains to note, technology is not an appropriate way to tackle bee-related issues (e.g. Robo-Bees [Gleadow et al., 2019])! In the case of hive health monitoring specifically, however, we feel that sensor-based approaches have considerable benefits that we explore in the remainder of this article.

## 2 Method

We examined the literature on hive monitoring sensor technology for any study of social hive bees, including honey bees, bumble bees, and stingless bees, that used electronic or electro-mechanical sensors at or near the hive to gather information about the insects, their health, behaviour, hive conditions, or interactions of their colony with its environment. We commenced our survey with the articles included in the review of Meikle and Holst (2015) [Meikle and Holst, 2015]. This provides a comprehensive survey up to 2015. We analysed articles listed in the survey [Marchal et al., 2019] covering work up to 2019. To identify more recent articles we used “Google Scholar” ^1^, querying keywords (“bee” OR “stingless bee” OR “honey bee”) AND (“monitoring” OR “sensor”). We also conducted reverse snowballing by examining the bibliographies of articles that came to light using these searches.

We constructed specific queries for stingless bees as just noted, since within the publications covered by Meikle and Holst (2015) [Meikle and Holst, 2015] and Marchal et al. (2019) [Marchal et al., 2019] work focused predominantly on other species.

We excluded studies that didn’t use physical sensors in their experimental methods, such as research on pre-existing datasets obtained independently using relevant sensors. Also, although sensor-assisted field monitoring of insects is important for insect pollination and biodiversity studies [Ratnayake et al., 2021b, Ratnayake et al., 2021a], and its approaches may overlap with hive monitoring, we chose to exclude such research from our survey to maintain our focus on hives.

After applying the above criteria to the literature, 147 articles in total remained for our study. These were published across a range of venues, but primarily peer reviewed conferences and journals. In the next section (sec. 3), we provide a detailed analysis of the articles, their contents and a synthesis of our findings.

## 3 Results

Here we propose a new, practical, structured framework for beehive monitoring and its associated technology Sec. 3.1. In Sec. 3.2, we illustrate the findings of our review by reference to this new framework. As we demonstrate below, a large part of the rationale for commercial hive monitoring products is to enable beekeepers to assess hive health without interruption to healthy hives (minimise hive inspections with permanent internal sensors), reducing the need for travel (minimise travel via remote monitoring), to provide early warning (minimise notification delay), and to provide accurate health assessments (minimise false positive and false negative reports of unhealthy hives). Hence, where possible, the examples provided have been selected to highlight the claimed, actual and potential for hive monitoring to inform us about colony health.

### 3.1 A framework for operational, investigative and predictive hive monitoring

People who monitor bees include those among the bee behavioural, ecological and pollination research communities; the beekeeping industries; hobbyists; and agricultural industries dependent on bees for pollination. Although there are overlaps, the objectives and priorities of each stakeholder group, even each individual beekeeper, may differ. For instance, while an entomologist might want to know the relationship between hive temperature and colony swarming behaviour, commercial beekeepers and hobbyists may wish for early alerts of hive health issues or imminent swarming so that they can respond accordingly. On the other hand, commercial growers might not be interested in swarming at all as long as their crops are being properly pollinated and the hive remains healthy. Given the complexity of goals for people working with bees, we find it beneficial to propose a simple framework for understanding beehive monitoring. Our framework consists of three monitoring activity types:

#### Operational monitoring

the routine collection of hive indicator data to obtain a snapshot of a colony at any time. Operational monitoring neither draws any conclusion nor provides any interpretation of its monitored data; this job, such as to infer hive health, is left to the end-user.

#### Investigative monitoring

the collection of operational hive indicator data for comparison against historical or external data to discover or research general relationships and to build an understanding of insect and hive behaviours, indicators, or properties, such as the baseline parameter values for a healthy hive.

#### Predictive monitoring

the collection of operational hive indicator data for comparison against known trends and behaviours gleaned from Investigative monitoring, in order to predict future insect or hive behaviour, or to infer current hive aggregate properties that aren’t directly observable or measurable (e.g., in particular, “hive health”).

We show below how this framework provides a functional classification that formalises and simplifies the process of articulating the expected outcomes or goals of a monitoring system or program. It reduces confusion by describing monitoring systems by reference to the intended *application* of the system or program, rather than the user, since, as indicated above, different user groups may have overlapping goals. The framework also describes monitors independently of the technology that might be applied. This is important since the development of past, current and future technologies is largely tangential to the need to monitor bees. Focusing on current technology buzzwords or trends, such as “Artificial Intelligence” for instance, can sometimes obfuscate rather than clarify the value of a monitor. For this reason we have designed our framework to focus on classifying *what* the monitors do, rather than *how* they do it. Further detail on this is provided below. Fig. 1 shows the relationships between the monitoring approaches we describe and the data they generate or require.

**Figure 1:**
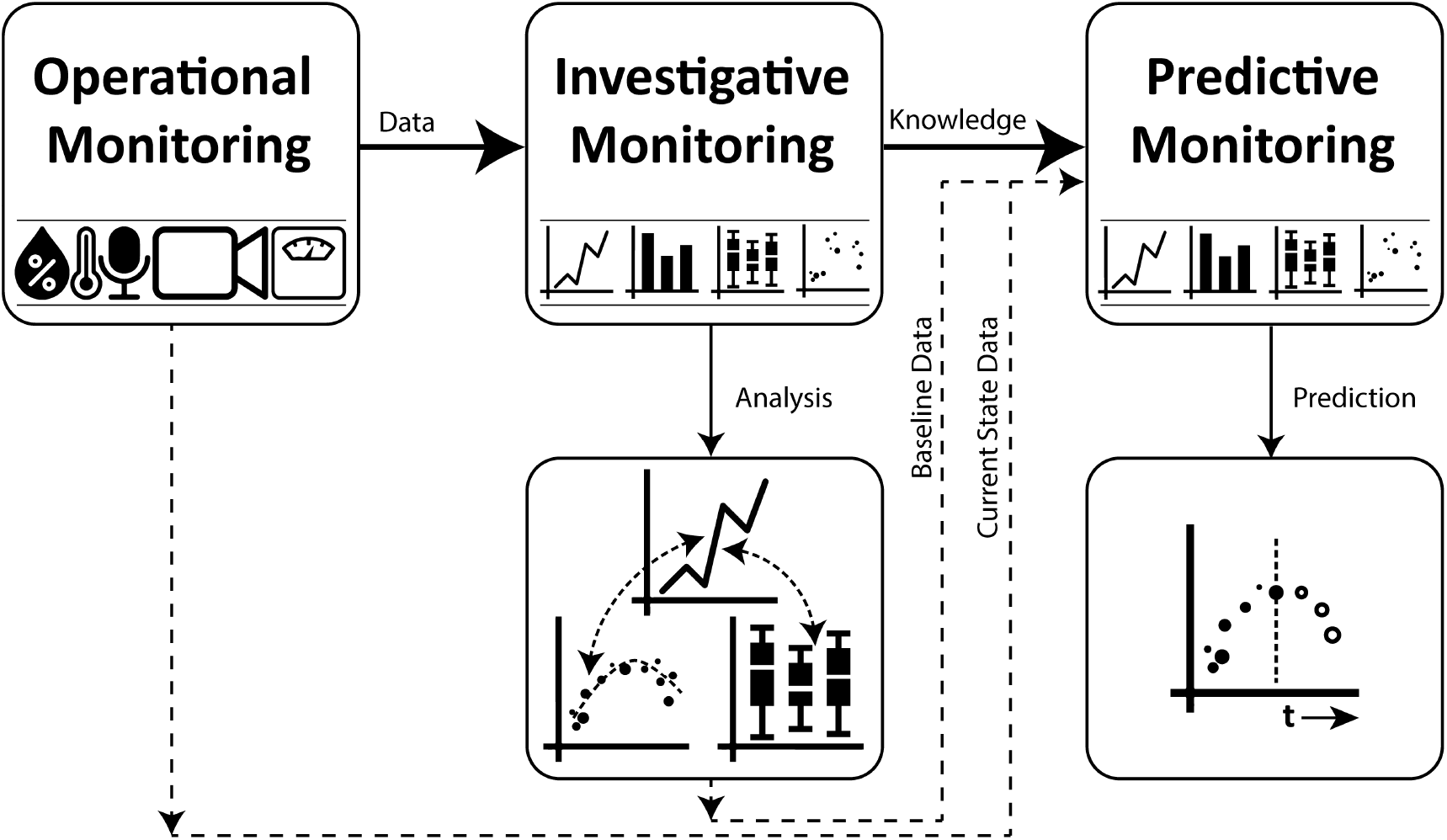
Relationship between **O**perational, **I**nvestigative, and **P**redictive monitoring in the OIP framework

Hereafter, we will refer to our framework as the “OIP” framework (Operational, Investigative, and Predictive).

This section provides detail on the OIP framework and the diverse ways in which technology is applied to support each type of monitoring.

#### 3.1.1 Operational Monitoring

Operational monitoring of beehives has recently gained appeal among the beekeeping industry and hobbyists, the evidence for this being the establishment of an industry dedicated to commercial products to monitor beehives (Table 1). The availability of these commercial products has made the digital monitoring process accessible to those with no electronics skills, and to those who do not have the capacity or dedication to devote to manual monitoring. Here, we discuss the hive parameters and diverse sensor technologies used for Operational monitoring.

**Table 1:** A few examples of commercial beehive monitoring products. As these are not research-specific products, none are marketed or specifically designed to perform Investigative monitoring. In practice, any Observational monitoring solution might be applied in this way. (R: Rechargeable, NR: Non-Rechargeable)

| Device, URL, health claim<sup>a</sup> | Sensors | Power Management | Connectivity | Monitoring Type |
| --- | --- | --- | --- | --- |
| Arnia<br>https://arnia.co/<br>“Better knowledge for bee health” | Audio, Temperature, Humidity, Weight, Light Sensor, Accelerometer, Bee Counter, Video | Battery (R), Solar Panel | 3G/4G | Operational, Predictive |
| ApisProtect<br>https://apisprotect.com/<br>“...identify which colonies need your attention...” | Temperature, Humidity, Audio, Accelerometer | Battery | - | Operational, Predictive |
| Buzzbox<br>https://osbeehives.com/<br>“...monitor the health of your colony from your smart phone” | Temperature, Humidity, Audio | Battery (R), Solar Panel | WiFi, Cellular | Operational |
| Broodminder<br>https://broodminder.com/<br>“...guides hobbyist beekeepers ...providing detailed insights about bees’ health” | Temperature, Humidity, Weight | Battery (NR) | WiFi, Cellular | Operational |
| Hive Mind<br>https://hivemind.nz/<br>“Remote beehive monitoring for higher yield and stronger hives” | Temperature, Humidity, Weight, Bee Counter, Rain Gauge | Battery (NR) | Satellite Hub | Operational |
| Hyper Hyve<br>https://hyperhyve.com/<br>“24/7 access to your hives data through a wireless connection to the cloud” | Temperature, Humidity, Weight | Solar Panel | Cellular | Operational |
| IOBee<br>https://io-bee.eu/<br>“Fighting honey-bee colony mortality through IoT” | Temperature, Humidity, Weight, Bee Counter | Battery | Cellular | Operational, Predictive |
| Pollenity<br>https://www.pollenity.com/<br>“...already helped save beehives, increase productivity...” | Temperature, Humidity, Weight, Acoustic | Battery (R) | WiFi | Operational, Predictive |

<sup>a</sup>All of the health claims collected from respective websites accessed on 11 Oct 2022.

##### Temperature and Humidity Sensors

Honey bees actively regulate temperature and humidity in their hives [Human et al., 2006,Southwick and Moritz, 1987] for healthy queen egg production, brood rearing, and food preservation. Therefore, access to these two hive parameters is fundamental to Operational hive monitoring. Often temperature and humidity are monitored together due to the availability of integrated sensors (i.e. SHT1X sensors) [Poposki and Gjorgjevikj, 2020, Kviesis et al., 2020, Cecchi et al., 2020, Cecchi et al., 2019, Terenzi et al., 2019, Ochoa et al., 2019]. In addition to being inexpensive, the simplicity and usefulness of temperature and humidity sensors makes them popular among stakeholders and integral to commercial products (see Table 1). Often, both internal and ambient temperature and humidity are recorded [Cecchi et al., 2020, Catania and Vallone, 2020].

##### Weight Sensors

Colony weight, one of the earliest hive indicators to be monitored [Hambleton, 1925, Gates, 1914], provides insight into hive health as a function of colony workforce and hive food stores. As Hambleton (1925) indicated [Hambleton, 1925], the Operational monitoring of colony weight was conducted even in the 1920s among the best beekeepers who maintained a “scale colony”. This hive was manually weighed to understand honey flow trends. The introduction of sensor technologies in the 1990s has rendered the process of continuous colony weight monitoring straightforward [L and C, 1990]. Recently, beehive weight scales capable of automatically sending periodic data to data storage using IoT (Internet of Things) and SMS technology with GSM protocols, have gained popularity for continuous monitoring [Hong et al., 2020, Poposki and Gjorgjevikj, 2020, Kviesis et al., 2020, Lyu et al., 2019, Zabasta et al., 2019, Ammar et al., 2019]. Many commercial products now include a scale to monitor hive weight, albeit some offer this only as an added service [Arnia, 2022, BeehiveMonitoring, 2022, HiveMind, 2022, Pollenity, 2022].

Scales are more expensive than temperature and humidity sensors and require slightly more elaborate setups. A useful weight scale’s range and resolution requirements for a hive can make them expensive. For instance, Kviesis et al. (2020) [Kviesis et al., 2020] proposed a system that includes a BOSCHE Wagetechnik Single point load cell H30A, rated for 200kg max load but costing 191.00 EUR/unit. Therefore, adding a weight scale to every hive of a professional beekeeper managing hundreds or thousands of them would incur a large upfront cost. On the other hand, since a large part of the expense comes from setting up the system to receive data from the scale, [Kviesis et al., 2020], if a hive is already rigged to other sensors, adding weight would be less expensive. Conversely, if a beekeeper first installs a weight scale, other sensors could piggyback on the electronics.

##### Audio, Video, and miscellaneous monitoring technologies

Audio, video, and other monitoring technologies such as gas sensors [Szczurek et al., 2019a], thermal imaging [Klein et al., 2014], infrared sensors [Shaw et al., 2010], GPS [Ntawuzumunsi et al., 2021], accelerometers [Kontogiannis, 2019], vibration detectors [Aumann et al., 2021], and light intensity sensors [Keppner and Jarau, 2016] are some less standard technologies we identified for Operational monitoring. Although they provide valuable data, perhaps their cost and complexity inhibits widespread adoption. The data gathered from these sensors are often high dimensional and can provide a good basis for subsequent Investigative and Predictive monitoring. For instance, from independent Operational monitoring of hive entrance activity, it is non-trivial to understand hive strength. However, with subsequent Investigative and Predictive monitoring initiatives, one could infer colony foraging activity or strength at any time [Ngo et al., 2019].

The sensor technologies just outlined underpin the Investigative and Predictive monitoring schemes discussed in the following sections.

#### 3.1.2 Investigative Monitoring

Investigative monitoring is the foundation of our understanding of bees. In its manual form, it dates back to Aristotle [Carlson, 2015] whose study of bees and their behaviour a few hundred years B.C.E. was largely unrivalled until the 17th century. However, this paper limits its discussion to recent Investigative monitoring approaches that rely on sensor technology.

Investigative monitoring is valuable to entomologists, ecologists, and behavioural scientists for its own sake as it provides the foundation for their sciences. It is also valuable as a precursor to Predictive monitoring (discussed later) since it can form the baseline against which predictions are made. We subdivide Investigative monitoring into three areas of interest below.

##### Understanding Bee Behaviour

Understanding bee behaviours such as swarming, absconding, clustering, fanning, and foraging is a strong driver for Investigative monitoring since understanding these phenomena has tangible practical value for bee pollination management, research value for entomologists, and conservation benefits for those concerned with biodiversity maintenance and sustainability of natural ecosystems.

###### Swarming and Absconding

*Swarming* is the process of honey bee colony reproduction where an old queen and about 30% to 70% of her workforce suddenly leave the hive for a new nesting place [Seeley et al., 1989]. Since this drastically depletes the hive workforce, understanding which hive indicators are related to swarming is helpful for beekeepers, especially if they later use this knowledge for decision-making informed by Predictive monitoring. It has been discovered that temperature increase inside the hive is a key indicator of imminent swarming [Kviesis and Zacepins, 2016]. Hong et al. (2020) [Hong et al., 2020] have determined the specific location inside the honey bee hive (*Apis Mellifera*) to monitor for this. Additionally, investigation of the sounds emitted by honey bee hives has proven effective for predicting swarming events [Chen et al., 2020, Zgank, 2019, Rybin et al., 2017]. Recent approaches to detecting this event include the combination of temperature, sound, hive weight, and humidity [Cecchi et al., 2020, Hong et al., 2020, Anand et al., 2018]. Researchers have investigated too how vibration [Aumann et al., 2021, Ramsey et al., 2020] and radar sensors [Aumann et al., 2021] can be used to detect swarming events.

*Absconding* is the process of the whole colony leaving a hive to find a new place to inhabit in response to stress or the unsuitability of their current situation. Absconding is more commonplace in the tropics than in temperate zones for honey bee species *A. cerana, A. florea, A. andreniformis, A. dorsta, A. laboriosa*, and African *A. mellifera* [Hepburn, 2010]. Kridi et al. (2016 and 2014) [Kridi et al., 2016, Kridi et al., 2014] have investigated how hive temperature data relates to whether or not absconding conditions will be met. From this work, it can be inferred how temperature data can be used to predict absconding.

###### Foraging

Foraging behaviour directly influences colony productivity and insect pollination capability [Ngo et al., 2021a]. It forms a key part of research on insect behaviour, as well as plant reproductive biology and evolutionary ecology. Therefore, understanding the foraging behaviour of bees is crucial to understanding their relationships with wild flowering plants, trees and horticultural crops [Zych et al., 2013], as well as a hive’s ability to produce honey, and in general, a hive’s interaction with its environment [Abou-Shaara, 2014]. The hive entrance is convenient for monitoring bee departures and arrivals on foraging trips [Ngo et al., 2019]. Since worker bees may disperse over many square kilometres during foraging bouts, the analysis of data collected at the hive entrance can sometimes be the only practical way to investigate their foraging activities. For instance, the mean distance travelled by honey bee foragers is 5.5km [Beekman and Ratnieks, 2000] while *Tetragonula carbonaria*, a stingless bee species native to Australia, can travel around 700m for foraging [Smith et al., 2016]. The extensive ranges of these fast, tiny insects present many difficulties for some kinds of in-field investigation (e.g. see [Ratnayake et al., 2021a, Ratnayake et al., 2021b]).

There are several approaches for monitoring foraging activity at the hive entrance. Photosensors placed in specially constructed entrance channels for honey bees to pass through [Erickson et al., 1975] provide an early example of a bee-counting sensor. More recently, constructing a bee-passage using infrared sensors [Anuar et al., 2021, Pesovic et al., 2017, JERRY et al., 2005] for bee counting at the hive entrance has been reported. Attaching RFID antennas at the hive entrance is also documented [Ai and Takahashi, 2021, Susanto et al., 2018] to monitor bee departures and arrivals where bees are fitted with small RFID tags. However, this approach is cumbersome and time-consuming as it requires the bees to be RFID-tagged in a labour-intensive process that is impractical for monitoring large workforces. Recently, continuous-wave radar [Aumann et al., 2021,Cunha et al., 2020, Woodgate et al., 2017] and LiDAR [Carlsten et al., 2011] have also been used for monitoring bee foraging activity. Additionally, Meikle et al. (2008) [Meikle et al., 2008] showed that within-day weight variation or “detrended weight” of a colony measured at the hive indicates foraging traffic variation.

##### Disease and Pest Control

Although pests and diseases have always been a threat to honey bee colonies, around 2006, the US beekeeping industry started reporting unusually severe bee losses, which contributed to the rapid collapse of whole colonies in a phenomenon dubbed Colony Collapse Disorder (CCD) [Evans and Chen, 2021]. Intentions to mitigate the potential impacts of such devastating pest and disease outbreaks, a direct threat to honey bee health with serious implications for food production in particular, justify much interest in honey bee hive monitoring. We have seen two approaches for sensor-assisted monitoring of honey bee hives for disease and pest control; image sensor-based approaches [Schurischuster and Kampel, 2020, Bjerge et al., 2019, Bauer et al., 2018, Schurischuster et al., 2018, Chazette et al., 2016, Elizondo et al., 2013] that include thermal imaging [Bauer et al., 2018], and gas sensor-based approaches [Bak et al., 2020, Szczurek et al., 2020, Szczurek et al., 2019a, Szczurek et al., 2019b]. Notably, almost all relevant research papers we examined confronted only the Varroa destructor mite [Bak et al., 2020,Szczurek et al., 2020,Szczurek et al., 2019a, Szczurek et al., 2019b], with an exception for detecting American Foul Brood [Bąk et al., 2022].

Image sensor-based approaches for detecting the presence of Varroa apply computer vision, machine learning and deep learning algorithms to detect mites from a video stream. Two ways this has been achieved are detecting infected cells inside the hive [Bauer et al., 2018, Elizondo et al., 2013] and detecting mites attached to bees as they pass through the entrance [Schurischuster and Kampel, 2020,Bjerge et al., 2019,Schurischuster et al., 2018,Chazette et al., 2016]. Both approaches require video capture (and possibly streaming) to be established beforehand. At the entrance, a tunnel can be built that restricts bees to a single-file passage, simplifying the image segmentation process and subsequent analysis for the presence of mites [Schurischuster and Kampel, 2020, Bjerge et al., 2019].

Gas sensor-based approaches use an array of gas sensors, an electronic nose (E-nose), to detect and classify complex odour patterns emitted from Varroa mites [Bak et al., 2020] and dead, rotting honey bee larvae infected by American Foul Brood [Bąk et al., 2022]. Since the chemical indicators of varroasis are unknown [Szczurek et al., 2019a], the authors of the Varroa detection method opted to use a sensor array instead of a single gas sensor. The output of this array was later passed through classification algorithms to detect and measure infestation levels. In the case of American Foul Brood detection, the authors used a similar technique where the measurements obtained from the gas sensor array were classified as indicative of either a sick or healthy colony.

##### Environmental & Ecological Monitoring

The use of honey bees as environmental and ecological bio-indicators is emerging as an approach to gather qualitative and quantitative environmental data by using the species’ predictable responses to environmental changes [Cunningham et al., 2022]. Honey bees can sense chemical signals and recognize vapour concentrations of as little as a few parts per trillion [Bromenshenk et al., 2015]. Additionally, they are very widespread and forage over long ranges (5.5km to 9.5km) [Beekman and Ratnieks, 2000], bringing pollen, nectar, resin, water, and other material back to a central hive. This makes honey bees excellent environmental bio-indicator species. Bromenshenk et al. (1985) used chemical analyses of honey bees collected from Puget Sound, Washington, USA to demonstrate the effectiveness of bees as large-scale environmental monitors [Bromenshenk et al., 1985]. Clearly then, the use of bees for monitoring is not new. More recently, bees have been used to monitor heavy metals such as Hg, Cr, Cd, and Pb in urban areas and natural reserves [Perugini et al., 2011], for sampling airborne Particulate Matter (PM) [Negri et al., 2015] to understand environmental pollution level, and for monitoring agricultural pesticide residue [Silvina et al., 2017]. Cunningham et al. (2022) [Cunningham et al., 2022] provides a broad review of honey bee biomonitors for environmental contaminants, plant and pollinator pathogens, climate change and antimicrobial resistance.

Other approaches to external environmental and ecological monitoring through honey bees include that of Zhao et al. (2021) [Zhao et al., 2021]. They investigated the use of a beehive’s sound emissions analysed using machine learning to detect air pollutants. Bayir and Albayrak (2016) [Bayir and Albayrak, 2016] utilized a wireless sensor network to measure the weight, humidity and temperature of control hives to determine the nectar flow period of a region.

The Investigative monitoring schemes discussed above provide useful data to inform entomologists, ecologists, and those studying complex social societies. As touched on already, many of them could potentially (or already do) serve as a baseline against which to make predictions about a monitored hive’s future state. We discuss this idea in more detail in the next section on Predictive hive monitoring.

#### 3.1.3 Predictive Monitoring

Insect behaviour prediction is a powerful application for hive monitoring that assists proportionate and immediate intervention crucial for hive maintenance. Timely alerts from Predictive monitoring systems raise beekeepers’ awareness of their hives’ needs and can enable them to ensure hives remain healthy whilst minimising the time they spend under stress. Predictive monitoring is especially useful if the hives are remote, or if a beekeeper has too many hives to inspect manually at regular intervals.

In the case of remote Predictive hive monitors, they must be connected to communications networks, such as via cellular phone towers or satellite (many hives are not within access of cabled network infrastructure), to enable them to communicate their predictions to users in a timely fashion – a late alert is history! In some remote areas, especially in large or developing countries where agricultural land is far from city infrastructure, this may be expensive or even impossible to achieve reliably. There is a spectrum across which connected Predictive hive monitors can therefore function with regard to their data transfer. At one extreme, the monitor is a dumb data logger that uploads its data to the cloud or other storage for processing, and then the processing software communicates its predictions to the user. Precision apiculture using IoT by Poposki and Gjorgjevikj (2020) [Poposki and Gjorgjevikj, 2020] is an example of this approach where the data logged by the system is sent to the web server which then processes it. In another example, Zhao et al. (2021) [Zhao et al., 2021] transmit the monitored hive audio signal to a cloud server where it is analysed for detecting air pollutants. This has the advantage of reducing the complexity, cost and power consumption of the remote devices, simplifying their construction and maintenance, as well as improving their battery life in cases where cabled power is inaccessible. At the other extreme, all data is processed at the edge, and only the subsequent alert is transmitted to the user. This requires a degree of computational power at the edge, which requires access to electrical power (perhaps a solar-charged battery for remote hives [Ammar et al., 2019]), and power management systems.

##### Swarming

Predicting swarming is an extremely time-sensitive application for Predictive monitoring since the time for the hive to warm up for swarm take-off is short (from 8 to 20 min) [Zacepins et al., 2016] for honey bees, and swarming duration is only 35*±*15 min [Ferrari et al., 2008]. As discussed in section 3.1.2, temperature, humidity, sound, vibration and radar sensors, as well as combinations of these, can be used for swarm prediction. However, due to the rapidity of the event, the time between the prediction and the onset of swarming is as important as prediction accuracy. While temperature-dependent schemes can predict a swarming event minutes before it happens [Zacepins et al., 2016], Ramsey et al. (2020) [Ramsey et al., 2020] showed that the vibration-spectra-based approach can predict swarming up to 30 days prior to the event, facilitating much earlier intervention by a beekeeper.

Although insect behaviours such as foraging, clustering, and fanning may potentially be important for us to predict in honey bees, this is currently not well investigated. Perhaps predicting these behaviours in advance does not yet clearly provide much value to beekeepers concerned with apiary management. Foraging predictions, we feel, are likely to be very important for pollination-dependent agricultural industries. Such information enables growers to manage the likely pollination coverage of colonies in advance. This helps them make early decisions about the number of hives required for a specific site, crop, and growing season. Therefore, we anticipate (and recommend) research be undertaken to predict future foraging behaviour. Along these lines, Clarke and Robert (2018) [Clarke and Robert, 2018] have used local weather conditions to propose predictive models of honey bee foraging activity. In our opinion, this could be extended to attempt to predict foraging patterns several days ahead.

##### Predicting Hive Health

Although we did not identify any consensus definition of hive health (see our discussion on Sec. 4.4), many studies tried to assess or predict hive health using measurable proxies. Understanding traffic at the hive entrance is a very commonly used proxy for hive health [Tashakkori et al., 2021, Cunha et al., 2020]. Internal hive condition such as temperature [Braga et al., 2020, Seritan et al., 2018, Meikle et al., 2017, Murphy et al., 2016], *CO*_2_ concentration [Seritan et al., 2018,Murphy et al., 2016], humidity [Paffhausen et al., 2021,Murphy et al., 2016], audio [Sharif et al., 2020,Pérez et al., 2016] and weight [Paffhausen et al., 2021,Hladun et al., 2016] have also been used extensively as proxies for hive health. Overwintering colony loss rate [Meikle et al., 2017,Becher et al., 2013, Ngo et al., 2021b] and pathogen infestation rate are also proxies for hive health [Jara et al., 2020]. As hive health assessment plays such a critical role in hive management, and underpins many developments in hive monitoring across Operational, Investigative and Predictive monitoring approaches, we devote a specific section (discussion Sec. 4.4) below to exploring this application.

### 3.2 Review Analysis

#### 3.2.1 Beehive Monitoring Classification

Figure 2 provides a high-level overview of the articles we considered in this study classified according to the OIP framework. Considering the data and information flow between the monitoring categories (Fig. 1), it is not unexpected to find that Operational monitoring provides the most common rationale, with 104 articles utilizing this approach in their research, either exclusively or in combination with other approaches. There were 70 and 19 articles on Investigative and Predictive monitoring respectively. The number of single-form applications reinforces the same finding with 47, 15, and 5 published articles utilizing exclusively Operational, Investigative, or Predictive monitoring approaches respectively. We found 10 articles utilizing all three monitoring approaches in their research. The other 47 articles utilized a combination of two different OIP monitoring classifications.

**Figure 2:**
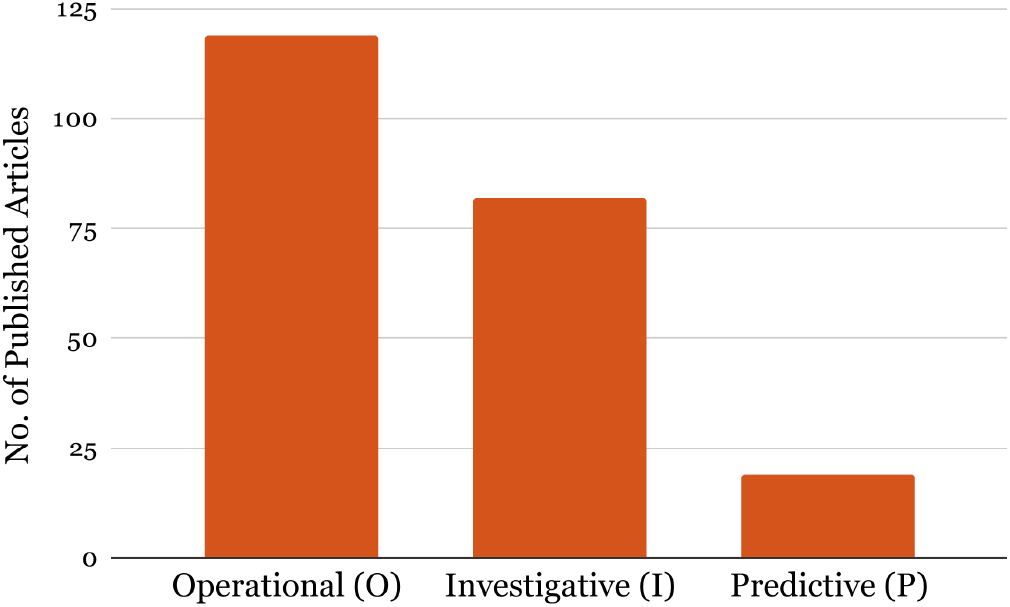
Classifying beehive monitors as Operational, Investigative, and Predictive (OIP) monitors

There may be several reasons why Predictive monitoring is currently relatively scarce. For prediction, continuous and automated monitoring of event-specific indicators is a prerequisite. For instance, to predict honey bee swarming events, there is a need to continuously monitor and analyse indicative proxy information that may be swarming-specific. Temperature or audio data might meet the requirements of this need if analysed in a specific way. Due to technological limitations, continuous and automated monitoring has only become feasible during the 90s [Meikle and Holst, 2015], and only become affordable much more recently. Secondly, predictions or information from Predictive monitoring must be delivered in a time-sensitive fashion if they are to be useful. For maximum value, predictive information ought to be transferred over a long range, since reducing beekeeper travel is, as noted, one major driver of this innovation and its adoption. Sending information over long distances from small, remote devices was only popularized recently with the introduction of the IoT for beehive monitoring. Consequently, when considered together, these reasons suggest why Predictive monitoring has only begun to establish itself as a monitoring approach since 2008. This is explored further in sec. 3.2.2.

#### 3.2.2 Beehive Monitoring Trends

Figure 3 clearly indicates that the number of publications about sensor-assisted beehive monitoring have trended positively over the last two decades, even though the idea of observing and trying to understand bees by making regular measurements is more than 100 years old. Another, perhaps unsurprising, finding is that Predictive monitoring is the youngest of three monitoring types, with publications from 2008 and numbers trending positively since then.

**Figure 3:**
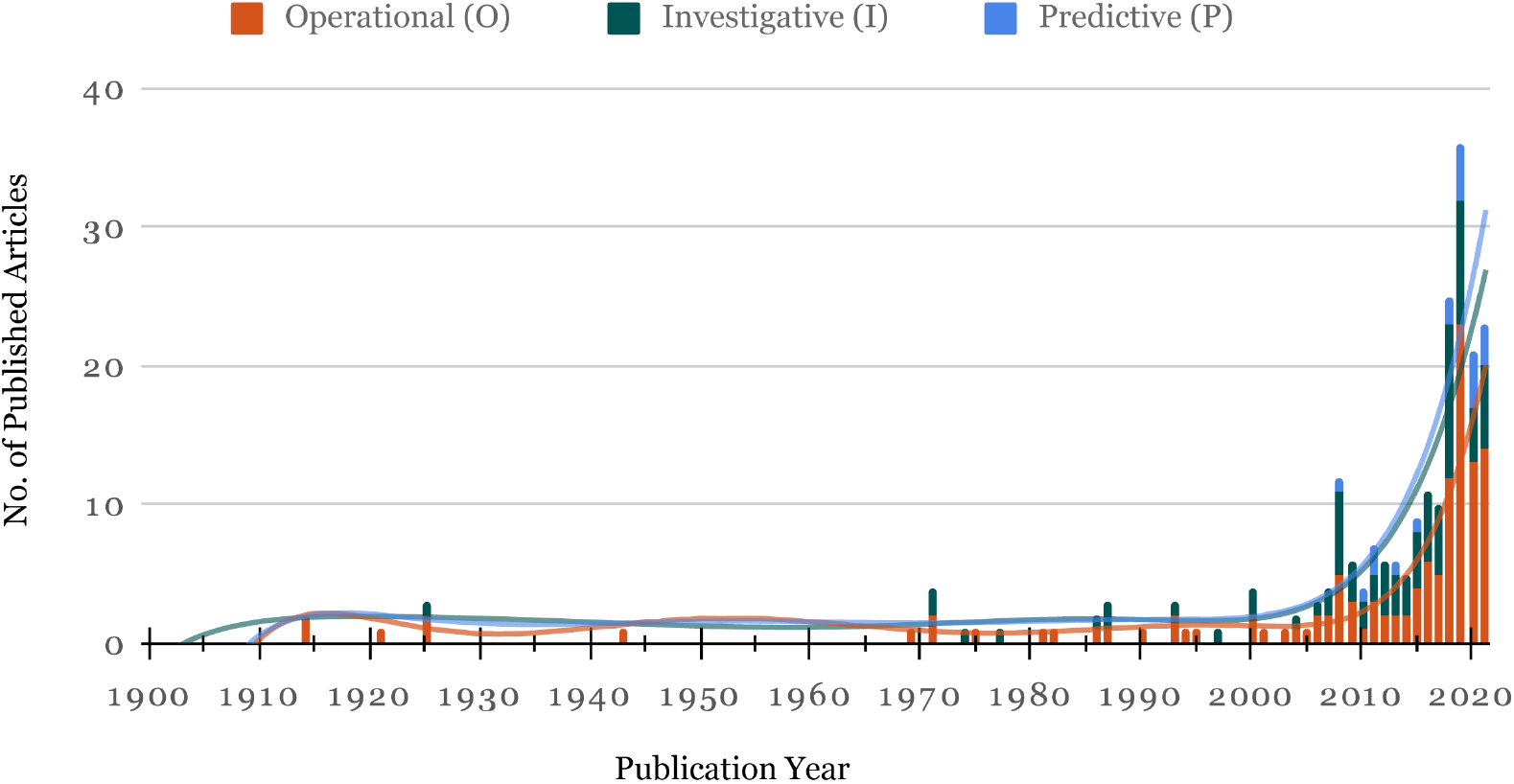
Trends in number of publications on beehive monitoring. The columns represent the number of published articles in each year. The lines represent trends for each of the monitoring types with *R*^2^ value of 0.777, 0.78, and 0.795 for Operational, Investigative, and Predictive monitoring respectively.

#### 3.2.3 Number of sensors used in monitoring projects

While the majority of the articles considered in this study utilized a single sensor for beehive monitoring, it was not uncommon to find multi-sensor approaches (Fig. 4a). We found evidence of up to 7 sensors used in a single project [Ntawuzumunsi et al., 2021, Ammar et al., 2019, Clarke and Robert, 2018]. It is also worth mentioning that, while the proportion of single and multiple sensor usage in Operational and Investigative monitoring is somewhat similar (45.19% and 57.14% single sensor usage for Operational and Investigative monitoring respectively), Predictive monitoring utilized a single sensor far more often than the multiple sensor approach (single sensors ware used 75% of the times in Predictive monitoring) as shown in Fig. 4b.

**Figure 4:**
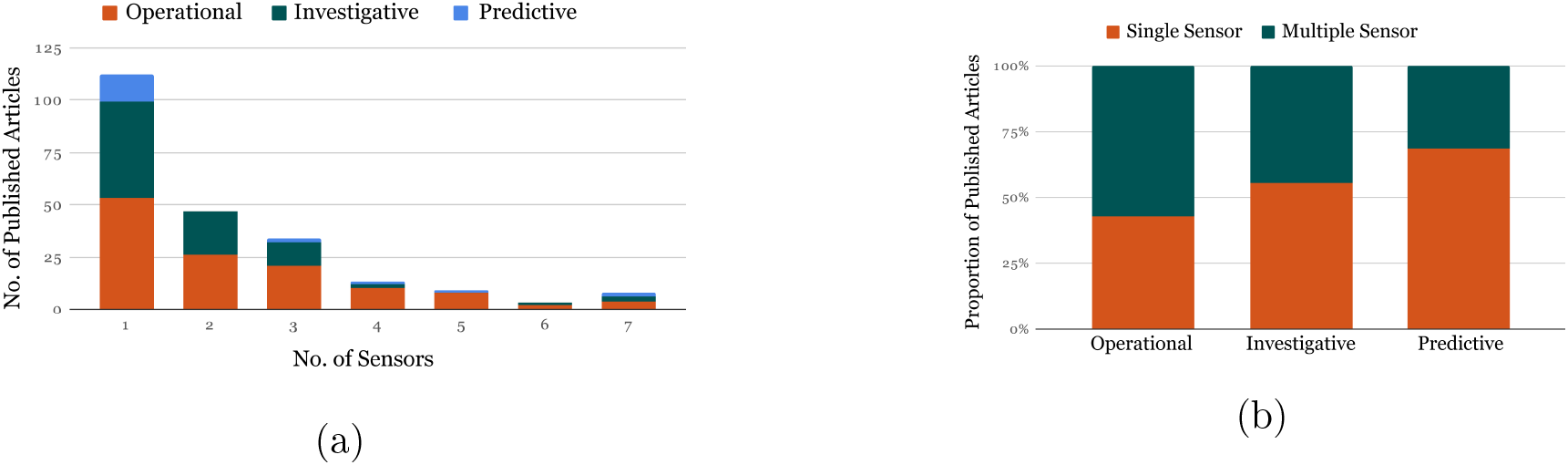
(a) Number of sensors included on a single hive monitoring system, (b) Use of single and multiple sensors across the three monitoring types

#### 3.2.4 Sensor Technology

While we found evidence that a range of sensor technologies are in use, the temperature sensor is by far the most common (21.13%, *n* = 60), with humidity (14.08%, *n* = 40) and weight (13.73%, *n* = 39) sensors closely following (Fig. 5). Other common sensors are image/video (8.10%, *n* = 23), audio (7.39%, *n* = 21), gas sensors (6.69%, *n* = 19), RFID (4.93%, *n* = 14), and bee counters (4.58%, *n* = 13). Temperature and humidity sensors were often combined as discussed in sec. 3.1.1. We also found that 15 of the sensor technologies were used across all OIP monitoring approaches, while 8 were shared between Operational and Investigative monitoring, 2 were shared between Operational and Predictive monitoring, and 2 were exclusively used for Operational monitoring. Fig. 6 demonstrates sensor technology shared across monitoring approaches.

**Figure 5:**
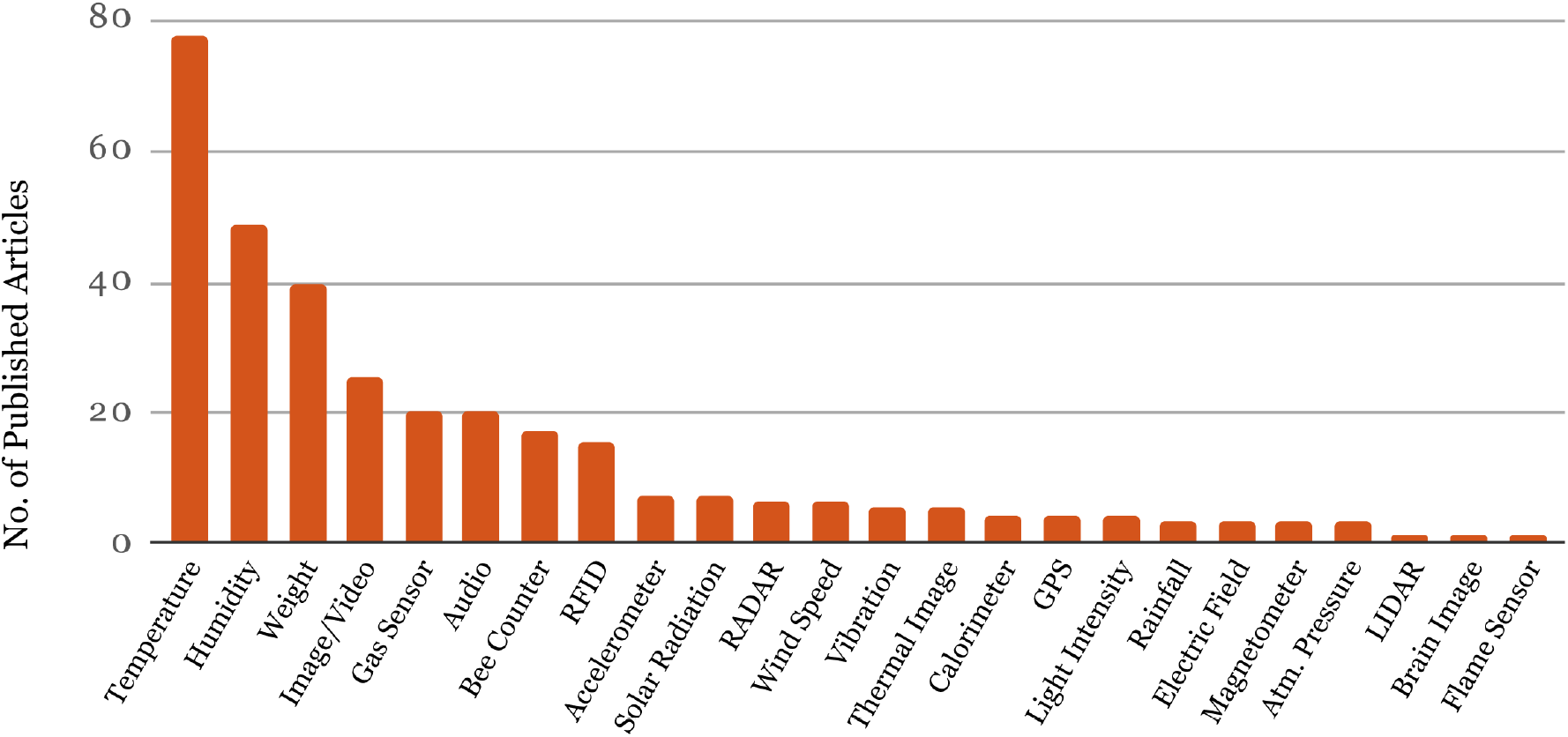
Ranked frequency distribution of sensor technology use in hive monitors

**Figure 6:**
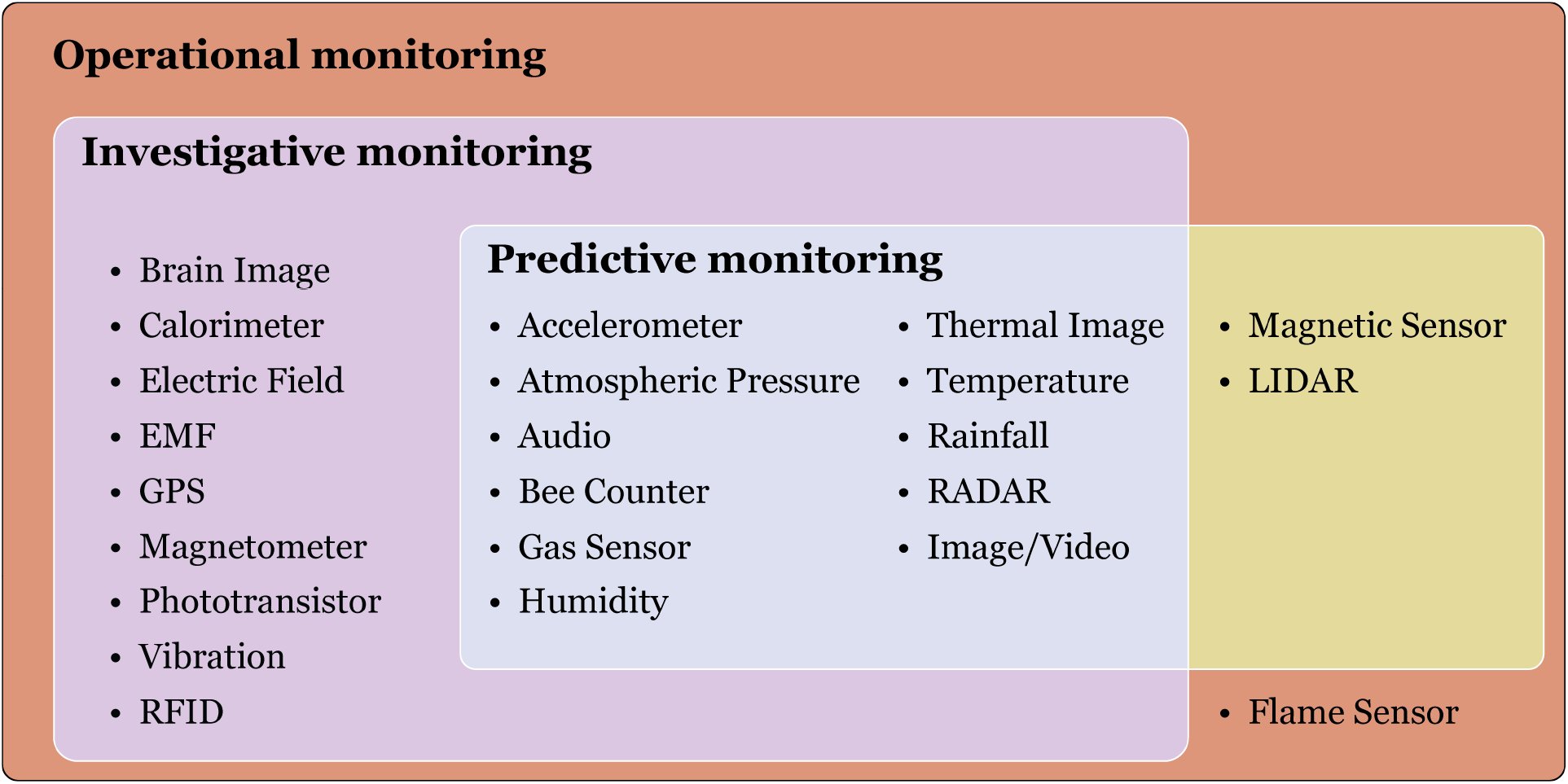
Venn diagram of the sensors used for different monitoring approaches. All of the sensors listed in this study have been used in Operational monitoring while a smaller subset have been used in Investigative and Predictive monitoring approaches both exclusively and in conjunction with other monitoring approaches.

#### 3.2.5 Sensor Connectivity

By sensor (or hive) connectivity, we are referring to how hive monitors and their sensors communicate data, or the outcomes of its analysis, to a processing server or the beekeeper. Connectivity requirements vary depending on the technology and monitoring type. Since Predictive monitoring is inherently time-sensitive, robust network connectivity is extremely important for this type of monitoring. Operational monitoring in most cases, and Investigative monitoring in some cases, also require reliable network connectivity. In the case of Operational monitoring, even though the technology itself may not predict hive health or other factors of importance to the beekeeper, if the beekeeper is to mentally infer these hive conditions from data in a timely way, then the data will still need to be reliably transmitted to them. If rapid response is not of concern, then the data generated from devices can be collected on-site and manually transferred off-site for subsequent analysis, eliminating the requirement for network connectivity. Possible networking choices include GSM [Seritan et al., 2018, Zacepins et al., 2017], Lora-WAN [Poposki and Gjorgjevikj, 2020, Zacepins et al., 2020], 3G [Kviesis et al., 2020, Murphy et al., 2016], ZigBee [Murphy et al., 2016], and MQTT [Tashakkori et al., 2021, Zabasta et al., 2019, Ochoa et al., 2019]. The choice depends on the monitoring technology, infrastructure availability and suitability. For instance, continuous, realtime transmission of audio and video requires high-bandwidth network connectivity (e.g. 3G/4G). On the other hand, temperature and humidity sensors can periodically transmit a tiny amount of data. Low power, low bandwidth networks such as Lora-WAN, ZigBee or MQTT, are probably well suited in this case.

#### 3.2.6 Geo-spatial Distribution

When referring to the location of the research we surveyed, we specify the location of any fieldwork described. When that wasn’t possible to determine, we used the location of the research institute of the article’s first author as the de-facto location of the study. If multiple locations were specified for fieldwork in a single study, we included them all in our analyses. Our mapping of study locations revealed that while the USA was the country with the highest number of research studies (28.38% of the total number, *n* = 42), among the continents, Europe had the highest number of studies (48.65% of the total, *n* = 72). Fig. 7 illustrates the distribution of hive monitoring research locations.

**Figure 7:**
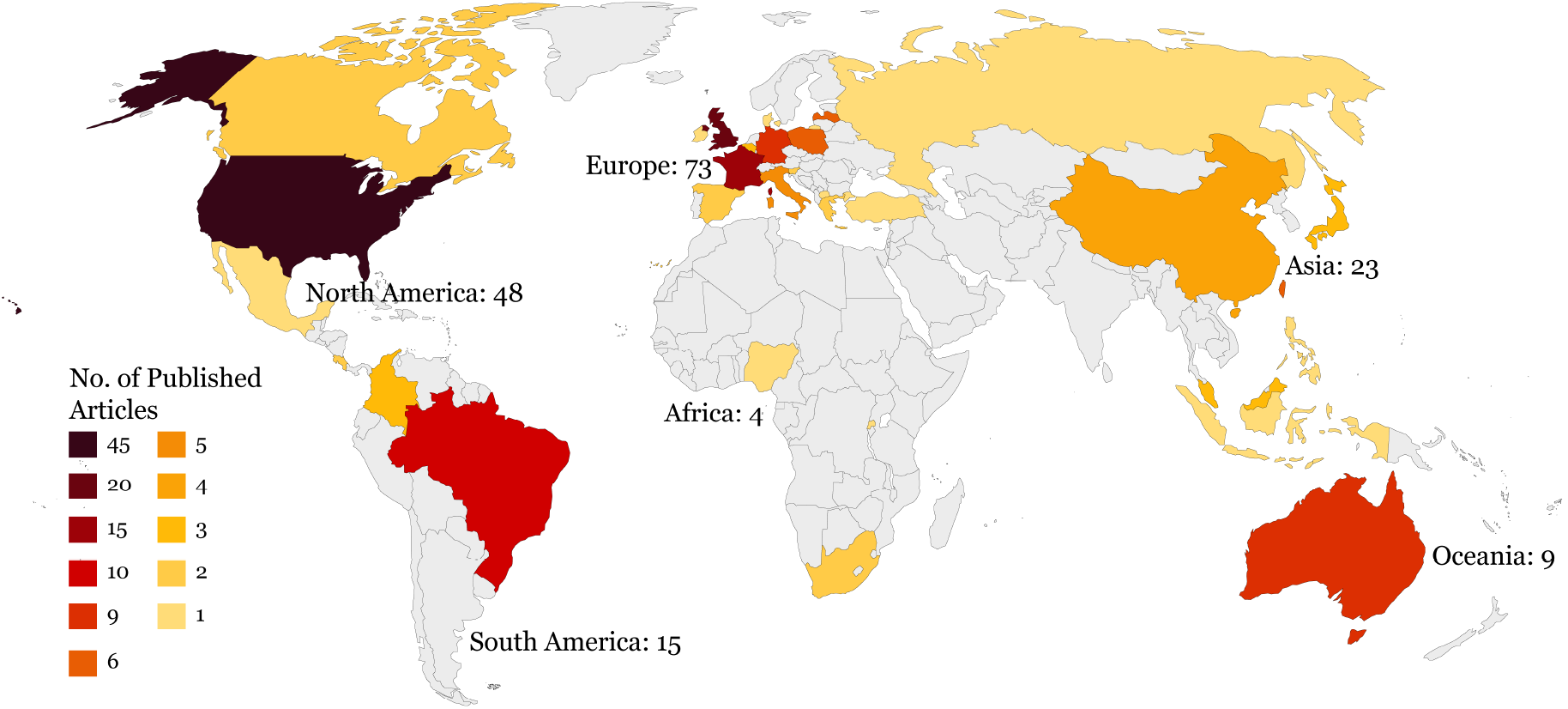
Distribution of beehive monitoring research locations.

#### 3.2.7 Insect Species Monitored

Several subspecies of European honey bee (*Apis mellifera*) were the dominant insects monitored by a large margin (Fig. 8), unsurprisingly given their widespread use as managed pollinators and honey producers. We also found evidence of Asian honey bee monitoring (*Apis cerana*) in some studies. These were exclusively performed in Asian countries. Bumble bees (*Apidae, Bombini*), and several species of Stingless bee (*Apidae, Meliponini*) were also monitored in some of the studies.

**Figure 8:**
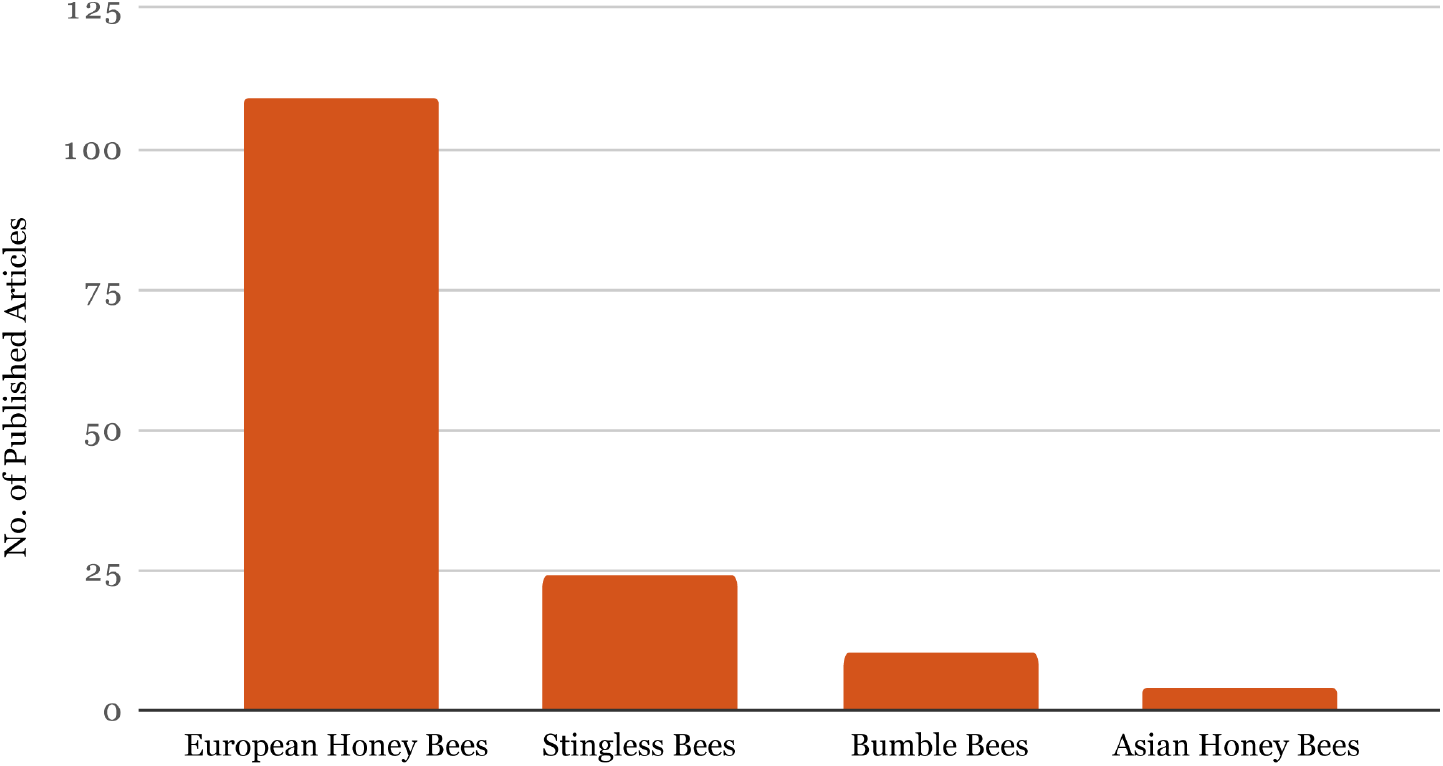
Bee varieties monitored

## 4 Discussion

To effectively monitor bees, automated, widespread, detailed sensor-assisted technology is required. An overwhelming majority (109 out of 147 articles) of the sensor-assisted monitoring of beehives we analysed was for European honey bees (Apis mellifera sp.). This is helpful for the honey bee and commercial pollination industries, but leaves other important species under-monitored. As bees continue to face a multitude of challenges, including climate change, environmental damage, loss of biodiversity, and disease and pests [Goulson et al., 2015, Gill et al., 2012, Kluser et al., 2010, Potts et al., 2010, Klein et al., 2007], the maintenance of this status quo leaves these insects, the crops they pollinate, and the natural ecosystems that depend on the services they provide, vulnerable – we simply lack the capacity to understand how the populations and behaviours of these key insects might be changing. In this section, we will discuss how sensor-assisted monitoring can help tackle challenges bees and bee habitats are facing, and in particular, how the understanding provided by our Operational, Investigative and Predictive monitoring framework can help us face challenges related to beehive health.

### 4.1 Climate Change

Climate change and the instability and natural disasters that result from it, such as wildfire, flood and temperature extremes, can have devastating impacts on bee populations [**?**]. Impacts are not always as simple as loss of hives; a recent study suggests that temperature extremes can cause a drop in sperm viability in honey bee queens [McAfee et al., 2020]. In the Australian state of New South Wales (NSW), the Apiarists’ Association (NSWAA) reported a loss of 5,000 hives during one freak event, the 2022 flooding [ABC-News, 2022]. However, this state has seen three extreme weather events in rapid succession: 2019-20 bushfires, 2021 mid-north coast flooding and 2022 northern flooding. In combination, these events are likely to have been even more damaging for bee species other than managed honey bees, especially for solitary native bees. But the extent of native bee losses are hard to determine since they are unmanaged and very difficult, expensive and time-consuming to monitor effectively. In fact, a substantial number of Australia’s native bee species have not even been properly identified, named or catalogued [Houston, 2018], let alone monitored. It is quite possible some as yet unidentified species have already been lost.

Sensor-assisted Operational and Predictive hive monitoring can provide service for fine-scale incidental environmental monitoring during natural disasters. Although local weather stations collect broad scale weather data, on-hive sensors can fill in some gaps whilst simultaneously allowing beekeepers to take preventive actions that reduce the damage caused to hives by extreme weather events including fires and floods. Sensor-assisted Operational and Predictive monitoring is also helpful for hives in remote forests and bushland. For instance, fire sensors on hives can provide early warning of natural fires and arson [Ntawuzumunsi et al., 2021].

### 4.2 Environmental Damage

Urbanization and agricultural deforestation have a profound impact on bees and other insects. Wild bees, in particular, are impacted by the loss and fragmentation of habitat, and floral resources [Olynyk et al., 2021, Kline and Joshi, 2020]. However, the impacts of environmental damage on the bee species have not been evaluated comprehensively [Olynyk et al., 2021, Kline and Joshi, 2020]. Evaluating environmental damage is challenging. Effective sensor-assisted monitoring can help reveal the extent of problems. For instance, Bayir and Albayrak (2016) [Bayir and Albayrak, 2016] determined the nectar flow period of a region by Operational monitoring of the temperature, humidity and weight of control hives using a wireless sensor network. The same approach could potentially be applied to evaluate a region’s recovery of floral resources following a wildfire. Although, if resources are too low, this may be detrimental to the health of a hive placed in situ as a biomonitor. Bees have been used as landscape-level biological monitors since 1974 [Bromenshenk et al., 1985] where environmental chemicals they collected from the air, soil, water, and vegetation were analysed. As Cunningham et al. (2022) [Cunningham et al., 2022] discuss, it is possible to monitor environmental contaminants such as heavy metals, persistent organic pollutants (POPs), particulate matters (PMs), agrochemical pesticides, and plant and pollinator pathogens by analysing the chemical and electronic composition of honey bee pollen, nectar, resin, and beeswax. However, since most of the processes discussed in the aforementioned article require sophisticated equipment (e.g atomic absorption spectrometry, scanning electron microscopy, X-ray spectroscopy, etc.), using these approaches on a large scale would be cost prohibitive. Alternative cost-effective approaches that can be installed cheaply on a large scale, possibly trading accuracy for cost, would be helpful. Gas sensor arrays used to detect Varroa destructor mites [Bak et al., 2020] and American Foul Brood [Bąk et al., 2022] are two examples of the cost-accuracy trade-off between sophisticated but potentially costly and time-consuming laboratory results, and rapid, cheap, but potentially less accurate field tests.

Unfortunately, the potential for bee species besides honey bees to inform us about environmental conditions is under-studied; in line with our findings that non-honey bee species are less studied in general. Investigations using sensor-assisted monitoring into how stingless bee hives respond to nectar flow may potentially be very useful, especially in areas where introducing honey bees is inappropriate.

### 4.3 Biodiversity Loss

A recent study suggests a global decline in bee species richness [Zattara and Aizen, 2021]. Evidence also suggests that the domesticated stock of honey bees is growing more slowly than the global pollination demands of agriculture [Aizen and Harder, 2009]. Whilst it may seem appropriate to recommend an increase in global domesticated honey bee stock to meet demand, relying so heavily on one species of managed pollinator is unwise. Honey bees are good pollinators for many crops [Hung et al., 2018], but there are many that honey bees do not pollinate well or at all [Garibaldi et al., 2016]. Also, a pest or pathogen outbreak (e.g. Varroa destructor mite) can devastate food security if we rely only on a single vulnerable bee species. For a robust pollination situation, we need diverse options with multiple levels of redundancy. In addition, wildflowers and native flowering plants depend on diverse bee species for pollination [Ollerton, 2016]. An increase of domesticated honey bee stock pressures other species, and consequently landscape biodiversity [Prendergast and Ollerton, 2022] and the robustness of pollination associated with it.

The need for robustness and diversity makes it important to study non-honey bees, such as stingless bees, that could be managed for pollination [Heard, 1999]. Operational and Investigative monitoring of stingless bees may help here. When extending current (predominantly) honey bee monitoring efforts to other species, especially to monitor wild social and solitary bees, our current approaches might not work. We may need to investigate a very different set of parameters than for honey bee hives. For instance, predicting swarming is a motivator for sensor-assisted honey bee hive monitoring, but is not helpful for stingless bees [Grüter, 2020b]. Therefore, we will need to adjust our approaches and technologies to target new colony phenomena and behaviours specific to new focal species.

### 4.4 Hive Health

A healthy colony is clearly integral to the well-being of any social bee. Hence, understanding hive health is of paramount importance to everyone dependent on managed pollination, bee products, or otherwise committed to the maintenance of hive bees. Consequently, it shouldn’t come as a surprise to discover that many of the monitors described above were developed as hive health monitors, as reflected in the “catch phrases” documented in Table. 1. In our survey of the 148 articles on monitoring, we identified 29 publications that discussed hive health in terms of hive properties. Sets of parameters were often considered as an indicative ensemble. Individually, these included:

#### Brood size and/or dynamics

Brood chamber volume [Greco et al., 2015] (for Stingless bees); brood production and survival [Hladun et al., 2016]; capped brood per colony [Jara et al., 2020].

#### Forager workforce size and/or dynamics

Population loss rate [Ngo et al., 2021b]; activity at the hive entrance [Paffhausen et al., 2021]; size of the overwintering colony [Becher et al., 2013]; traffic at the hive entrance [Tashakkori et al., 2021]; bee behaviour at the hive entrance [Schurischuster et al., 2018]; overwintering loss rate [Meikle et al., 2017].

#### Hive internal environment

Temperature, humidity, weight [Paffhausen et al., 2021]; temperature and *CO*_2_ concentration [Seritan et al., 2018]; thermal homeostasis [Meikle et al., 2017]; [Szczurek et al., 2019b] provides a list of citations on “physical and chemical conditions” including temperature, humidity, appearance, and weight of the hive as well as notes on gas content, valeric acid, caprylic acid, isocaprylic acid in the context of American foulbrood detection cited from Gochnauer and Shearer (1981) [Gochnauer and Shearer, 1981]; internal temperature and beehive weight [Braga et al., 2020]; humidity, temperature and *CO*_2_ [Murphy et al., 2016].

#### Hive audio

A variety of audio properties of hives were considered by [Terenzi et al., 2020, Sharif et al., 2020, Pérez et al., 2016], and mentioned in [Szczurek et al., 2019b].

#### Hive resources

Wax, honey [Hladun et al., 2016] (Potentially, any weight monitor could come under this category once hive biomass variation is accounted for).

#### Pathogen infestation

Varroa destructor, Nosema parasites, Deformed Wing Virus load [Jara et al., 2020] (Also see American foulbrood notes and references listed under Hive internal environment above.)

Notably, Braga et al. (2020) [Braga et al., 2020] collected six metrics, including the components listed above, to combine into a Health Score. Among the components is, uniquely, an assessment of stressors that might lead to reduced colony survival or growth potential (e.g. environmental factors external to the hive), and an assessment of whether the space is sanitary, defensible, and spacious enough for future egg laying. Independently of monitoring, to indicate the diversity of perspectives, the health of a hive has also been interpreted as a function of microbial balance among bees and colony [Anderson et al., 2011].

Adding to the heterogeneity of methods to assess hive health we identified in the literature, none of the surveyed papers provided a clear, formal, definition of hive or colony health, only a list of indicators. There certainly was no explicitly shared understanding of the concept. This is worrying as underlying assumptions about what should be measured, how it should be measured, why it should be measured, and how to interpret the meaning of particular measurements, are therefore being made implicitly, without agreement on the underlying concept to be inferred. Clearly, agreement on the “target” is essential for coherent monitoring technology design, implementation and application. This would seem to be a basic and important discussion to have with an aim to formalise agreement on how to direct (or assess) hive monitoring technology’s value for health monitoring.

One obvious drawback of the current ad hoc approach to defining hive health within the community of researchers and developers of monitoring technology, is that a subset of hive indicators must always be selected for attention, and others must be omitted, all without overarching guidance as to priority or appropriateness. This may lead to misapplication of a device, or suboptimal designs for specific use cases.

In a particular scenario, the selected hive parameters may be partially or completely irrelevant, or at least redundant, adding unnecessarily to the cost, complexity and perhaps usability of the technology, without necessarily benefiting users or bees. To take an extreme example, a hive monitored at its entrance for bees carrying Varroa destructor might be nearing collapse for countless reasons while the monitor reports “Clear of Varroa”. The hive may even be completely dead! Obviously this case can be avoided by common sense, but more subtle errors could occur in many ways depending on which parameters are monitored. It is also likely that a parameter’s relevance (not just its healthy value range) may vary between bee species. For instance, monitoring for the loss of a stingless bee hive’s queen may be less valuable as an indicator of concern than in the case of a honey bee hive, since stingless bees naturally create more queens [Grüter, 2020a].

It is beyond the scope of this article to formalise the definition of social insect colony health, although see [Braga et al., 2020]. Nevertheless, it is worthwhile to note here that hive health is a holistic property independent of our ability to monitor it. A healthy hive should be defined as a combination of past, present and foreseeable future conditions enabling it to maintain and reproduce itself. Such a definition therefore must include the degree of freedom from, or suppression of, toxins, diseases, pathogens, pests and other infestations. It should account for the interactions of the hive with its surroundings too since no hive can be considered healthy unless it is situated within a region of suitable micro and macro-climate, and sufficient diversity and quality of resources. Lastly, the current snapshot of internal state must be appropriate for a specific hive, in a specific environment, at a specific time. We feel that “appropriateness” should be understood here in terms of a hive’s ability to maintain and reproduce itself, and its robustness to likely perturbation and environmental variation. This is preferable to considering appropriateness in terms of crop pollination value, or saleable honey-production capability. After all, the hive is a biological and ecological entity in its own right, not simply a commodity for exploitation.

## 5 Conclusion

In this study, we have explored existing sensor-assisted beehive monitoring and established a framework facilitating adaptation and innovation of its technology. We have proposed a classification for hive monitoring systems and approaches, consisting of Operational, Investigative, and Predictive (OIP) monitoring, and have analysed previous research in terms of this. The proposed framework newly frames monitoring research in terms of its rationale, rather than relative to the technology, parameters, or hive conditions being monitored or inferred.

Quantitative manual Operational hive monitoring is over a hundred years old. Consequently, sensor-assisted Operational monitoring that automates its processes dominates monitoring research. Investigative monitoring has subsequently seen a steady increase since 2003. Predictive monitoring, however, is relatively new, commencing in 2008 and still only sporadically conducted. Predictive monitoring appears nevertheless set to become the new frontier in sensor-assisted beehive monitoring. Some current commercial hive monitoring products are already offering such capabilities, and recent advances in machine learning on small, portable, low-powered devices, suggest imminent expansion of in situ Predictive monitoring that was previously infeasible.

Our proposed OIP framework has enabled identification of shortcomings and gaps in hive monitoring research and technology. Compared to European honey bees, other hive bees such as Bumble bees, Stingless bees and Asian honey bees, are under-represented across the three monitoring classes. Since Investigative and Predictive monitoring rely on Operational monitoring as their data source, and Predictive monitoring requires knowledge and baseline analytical data from Investigative monitoring, the gap in Operational monitoring for these bees is consequential – it must be filled before meaningful Investigative and Predictive monitoring can be performed for them. Our analysis also revealed a disparity in geo-spatial distribution of hive monitoring. Unsurprisingly, developed nations are way ahead of the developing nations, yet developing nations stand to gain considerably from the benefits of sensor assisted monitoring. On the other hand, since we know the relationship between Operational, Investigative, and Predictive monitoring, developing nations can piggyback on the knowledge gained from years of Operational and Investigative monitoring in developed nations to start Predictive monitoring for well-established phenomena. This can help redistribute monitoring efforts to regions where honey bee pollination is problematic, where bee health is of particular concern, or where bees are particularly important to maintain tenuous food security. Of course, diligence is required to properly utilise all three types of monitoring if a phenomenon or environmental condition of importance is regional or sub-species specific. But inevitably, it is in regions of the greatest disadvantage, that careful, targeted sensor-based hive monitoring is most likely to be of the greatest benefit to people, ecosystems and bees alike.

## Footnotes

1 https://scholar.google.com

